# *Promfusion*: a synthetic fusion promoter enabling enhanced and balanced photoreceptor transgene expression

**DOI:** 10.64898/2026.06.05.730342

**Authors:** Sophie Tran, Jeanne Trinquier, Timothe Van Meter, Emilia A Zin, Céline Nanteau, Luisa Riancho, Anais Potey, Amélie Slembrouck-Brec, Maëlle Delmas, Ulisse Ferrari, Olivier Goureau, Deniz Dalkara

**Affiliations:** Sorbonne Université, INSERM, CNRS, Institut de la Vision, F-75012 Paris, France

**Keywords:** AAV gene therapy, Photoreceptors, Synthetic promoters, Retinal organoids

## Abstract

Achieving efficient and balanced transgene expression in both rods and cones remains a major challenge in retinal gene therapy. Current promoters either lack specificity or fail to provide sufficient cellular coverage and expression level. To address this limitation, we developed and evaluated two fusion promoters, Pikali and Nocchu, by combining PR1.7, a cone-specific promoter and GRK1, a promoter most active in rods. Here, we show that Pikali and Nocchu outperform their parental promoters, driving broader and more balanced GFP expression in rods and cones of human iPSC-derived retinal organoids. These constructs achieved transduction in 30% to 45% of photoreceptors, with higher expression levels than GRK1 and broader cellular coverage than PR1.7. Our findings establish Pikali and Nocchu as excellent candidates for retinal gene therapy, overcoming the limitations of existing promoters. By combining specificity, efficiency, and extensive photoreceptor targeting, these fusion constructs represent a novel and promising strategy for next-generation gene therapy vectors, addressing inherited retinal dystrophies and advancing clinical translation.

## INTRODUCTION

Promoters play a central role in gene expression by driving the transcription of specific genes in a controlled and cell-type-specific manner. In a genome, promoters are regulatory DNA sequences located upstream of genes that recruit RNA polymerase and other transcription factors to initiate transcription. Natural promoters have evolved to regulate gene expression with high specificity, but they are often limited by their size, cell-type restriction, or low activity outside their native context. These limitations pose significant challenges in gene therapy, where precise and efficient transgene expression in target cells is essential for therapeutic success and safety^1^. To overcome these challenges, researchers began modifying natural promoters by truncating them, enhancing key regulatory elements, or combining motifs from different sequences to improve activity and specificity. Over the years, synthetic promoters have been developed to address this need, often by deriving sequences from endogenous promoters^2^.

The PR1.7 and GRK1 promoters are important gene regulatory sequences used in retinal gene therapy to target specific cell types in the retina. PR1.7 is a 1.7 kb truncated version of the human L (red) cone opsin promoter (derived from PR2.1), developed for strong and specific cone photoreceptor gene expression. It was created by PCR amplification of relevant portions of the synthesized PR2.1 promoter and has been extensively tested in mice, dogs, and non-human primates ^3,4^. In non-human primates, it achieved robust and specific targeting of GFP expression in L/M and S cones. The PR1.7 promoter is highly efficient at directing expression in primate cones, showing robust GFP expression in L/M- and S-cones in non-human primates, dogs, and human retinal models, with minimal expression in rods and no leakage toward the inner retina^3,4^. However, it may exhibit species-specific differences in expression patterns. The GRK1 promoter, on the other hand, is used for rod-specific gene expression in retinal gene therapy^5,6^. It has been tested in canine retinas and other animal models, including both normal and mutated retinas. In normal canine retinas, it directs expression exclusively to rods, while in mutated retinas, it can drive expression in both rods and cones, which may be beneficial for certain disease conditions^7^. However, expression patterns may vary between species and between normal and mutated retinas, and it may not be truly cone-specific in all conditions. The development and testing of these promoters underscore the importance of careful evaluation across different species and disease states when developing gene therapy strategies for retinal diseases. The choice between these promoters depends on the specific therapeutic goals and target cell populations within the retina.

GRK1 shows variable specificity across species and disease states, directing expression exclusively to rods in normal canine retinas but potentially expressing in both rods and cones in mutated retinas^7^. This variability may limit its effectiveness in consistently targeting both photoreceptor types. Notably, GRK1 has been employed in clinical gene therapy approaches for Leber Congenital Amaurosis type 10^8^, where efficient expression in both rods and cones is considered important, further highlighting the importance of carefully selecting promoters to achieve appropriate photoreceptor targeting in a clinical context. The PR1.7 promoter, while efficient at directing expression in primate cones, shows minimal expression in rods, potentially reducing its effectiveness in therapies requiring expression in both cell types. Both promoters may exhibit species-specific differences in expression patterns, complicating translation from animal models to human therapies. In contrast, the CRX promoter offers advantages in targeting both photoreceptor types. CRX is essential for both rod and cone photoreceptor development and maturation, explaining the broader expression profile of its promoter in both cell types. While the CRX promoter appears to offer advantages in targeting both photoreceptor types, its 2.5kb variant^9^, though technically compatible with AAV vectors, leaves little space for therapeutic transgenes given AAV’s ∼4.7kb packaging limit. Although a smaller version has been recently reported in a single study, it currently lacks a complete characterization^10^. There is thus and unmet need for a smaller promoter sequence targeting both rods and cones, namely in both murine and human species for clinical application.

Current strategies for developing improved promoters have evolved significantly, leveraging both traditional and cutting-edge techniques. High-throughput screening of synthetic sequences, involves the systematic testing of libraries containing hundreds of synthetic promoter candidates^11^. These libraries are designed using a combination of strategies, including phylogenetically conserved DNA elements, known transcription factor binding sites, and epigenetic profiling, to maximize cell-type specificity and expression strength. Jüttner et al. developed and screened a library of 230 synthetic promoters across murine, non-human primate, and human retinal systems^11^. This extensive approach successfully identified synthetic promoters with specific activity in various retinal cell types, including photoreceptors. However, while the study demonstrated the potential of these promoters, the overall validation of promoters achieving dual and balanced rod-and-cone targeting was not extensively covered. Additionally, challenges such as ensuring robust activity across species, addressing variability in promoter performance between models, and achieving efficient and balanced targeting of both rods and cones remain unresolved. These findings highlight the need for further research to develop promoters that meet the stringent criteria required for therapeutic application.

More recently, artificial intelligence (AI) has been integrated into promoter design, providing a powerful tool to predict functional sequences based on known regulatory motifs and transcriptional patterns. AI algorithms can process vast amounts of genomic data to generate novel promoter sequences with the potential for high specificity and robust expression^12,13^. Despite its promise, this approach still faces key challenges. For instance, many AI-generated sequences require experimental validation to confirm their efficacy and cell-type specificity, as in silico predictions alone cannot account for all variables influencing promoter function in vivo. Moreover, the sheer number of possible promoter designs generated by these methods complicates the selection process, making it difficult to prioritize candidates for downstream testing.

These challenges underscore the limitations of current methodologies. High-throughput screening produces a wealth of potential candidates but requires significant resources for validation, while AI-based approaches, though innovative, are not yet enough to reliably predict de novo efficient promoters for human use. The need for improved strategies to streamline promoter discovery and validation remains a critical bottleneck in the field of gene therapy.

To address the challenges of achieving dual rod-and-cone targeting, we explored an approach based on fusing two complete and validated promoters. Promoters are composed of motifs that drive expression in specific cell types. By combining functional motifs from promoters optimized for rods and cones, we aim to create a single promoter capable of dual specificity and high efficiency. Unlike the widely used CAG promoter, which integrates specific enhancer and promoter elements (e.g., CMV-IE enhancer and β-actin promoter) to achieve broad expression^14^, our strategy directly combines synthetic promoters that have already been validated individually for rods and cones. This approach leverages their complementary strengths to create a single, efficient construct, addressing the limitations of existing promoters, such as insufficient coverage or low expression levels, and opening new avenues for efficient transgene delivery in retinal gene therapy.

To validate our hypothesis, we tested our fused promoters in two complementary models: the mouse retina, an established in vivo system for studying retinal transgene expression, and mature retinal organoids derived from human induced pluripotent stem cells (iPSCs), an in vitro model that closely mimics the genomic and transcriptomic landscape of the human retina^15,16^. While the mouse model provides a robust and accessible platform for in vivo testing, human organoids are indispensable for assessing promoter performance in a clinically relevant context. Indeed, the regulatory elements driving gene expression in mice often behave differently in human retinal cells due to species-specific differences in transcriptional landscape. By incorporating a human-relevant system into our study, we aim to ensure that the promoters we develop are not only effective but also specifically tailored to human biology, thereby addressing a critical barrier in the transition from preclinical research to therapeutic applications.

Our work not only seeks to develop and thoroughly characterize an efficient promoter for expression in both rods and cones but also introduces promoter fusion as a versatile strategy for designing new promoters. This proof of concept could pave the way for broader applications in gene therapy, extending beyond retinal diseases to any other genetic disorders requiring precise and cell-specific gene delivery.

## METHODS

### AAV production and titration

The plasmids used for AAV production were designed to include one of the four studied promoters upstream of the coding sequence for enhanced green fluorescent protein (EGFP), which served as a reporter gene. All plasmids were purchased from VectorBuilder. AAV vectors were produced a previously established protocol using the triple co-transfection method and the recombinant AAVs were purified via iodixanol gradient ultracentrifugation^17^. Vector stocks were quantified using quantitative PCR (qPCR) with SYBR Green (Thermo Fisher Scientific), employing primers specific to the AAV inverted terminal repeats (ITRs)^18^. Titrations were performed relative to a standard curve for precise determination of viral genome copies.

### Subretinal injection in mice

For this study, C57BL/6J wild-type mice (Janvier Laboratories) were used. Mice were anesthetized using isoflurane inhalation, and their pupils were dilated prior to injection.

Subretinal injections of 1 μL were performed under an operating microscope (Leica Microsystems, Ltd.) using a Hamilton syringe fitted with a 33-gauge blunt needle (World Precision Instruments, Inc.). Following the procedure, an ophthalmic ointment (Fradexam) was applied to protect the eyes. One month after injection, mice underwent fundus imaging under isoflurane anesthesia using a GFP-specific filter to assess transgene expression prior to euthanasia and enucleation.

### Generation of AAVS1::Arr3P H2BmCherry iPSC line

The generation of stable iPSC clones was performed as previously described^19^ using the co-transfection method with human codon-optimized SpCas9 and chimeric guide RNA expression plasmid and plasmid expressing Puromycin-resistant gene and H2B-mCherry under the control of partial mouse cone Arrestin (Arr3) promoter^3^ between AAVS1 homology arms. Correct integration of the Arr3P_ H2B-mCherry cassette was evaluated by PCR. After validation by sequencing one homozygous reporter cell line was selected, and pluripotency was assessed by immunostaining for pluripotency markers in iPSC colonies as well as markers of the three germ layers after trilineage differentiation (Fig.S2).

### iPSC Culture

Experiments were mainly conducted with two established human iPSC lines, iPSC-5f^20^ and the fluorescent reporter AAVS1::CrxP_H2BmCherry-iPSC line derived from iPSC-5f^19^.

Another fluorescent reporter iPSC line AAVS1::Arr3P_H2BmCherry-iPSC (Fig.S2) has also been used for specific experiments. Human iPSC lines were maintained on plates coated with vitronectin in mTeSR Plus medium (StemCell Technologies) at 37 °C in a 5% CO₂/95% air atmosphere. The medium was refreshed every 2 days. Cells were passaged weekly using Gentle Cell Dissociation Reagent (StemCell Technologies) at room temperature for 6 minutes. Detached cell aggregates were collected, gently dissociated into uniform suspensions, and replated onto fresh vitronectin-coated plates.

### Retinal Differentiation of iPSCs

Retinal differentiation was performed following a previously established protocol^20,21^. At approximately 70–80% confluence, iPSC colonies were transitioned into TeSR-E6 medium (StemCell Technologies) to initiate differentiation (Day 0). On Day 1, the medium was refreshed with fresh TeSR-E6 medium to minimize cell death. On Day 2, the medium was replaced with E6N2 medium, consisting of TeSR-E6 medium supplemented with 1% N2 Supplement-A (StemCell Technologies), 10 U/mL penicillin, and 10 µg/mL streptomycin (Thermo Fisher Scientific). Medium changes were performed every 2–3 days.

On Day 28, emerging retinal organoids were manually isolated using a needle and cultured as floating structures in ultra-low attachment 24-well plates (Corning). The organoids were maintained in ProB27 medium supplemented with 10 ng/mL recombinant human FGF2 (PeproTech). ProB27 medium consisted of DMEM:F12 (1:1, L-glutamine, Thermo Fisher Scientific), 1% MEM non-essential amino acids (Thermo Fisher Scientific), 2% SM1 Supplement (StemCell Technologies), 10 U/mL penicillin, and 10 µg/mL streptomycin (Thermo Fisher Scientific). Half of the medium was changed every 2–3 days.

On Day 35, retinal organoids were cultured in DMEM:F12 (1:1, L-glutamine, Thermo Fisher Scientific), 1% MEM non-essential amino acids, 2% SM1 Supplement, supplemented with 10% FBS (Thermo Fisher Scientific), 1 mM Glutamax (Thermo Fisher Scientific), 40 U/mL penicillin, 40 µg/mL streptomycin, and 1 µg/mL amphotericin B (Thermo Fisher Scientific). For photoreceptor maturation studies, organoids were transferred on Day 84 to DMEM:F12 (1:1, L-glutamine), 1% MEM non-essential amino acids, containing 2% B27 Supplement without vitamin A (Thermo Fisher Scientific) instead of SM1 supplement and supplemented with 10% FBS, 1 mM Glutamax, 40 U/mL penicillin, 40 µg/mL streptomycin, and 1 µg/mL amphotericin B. This maturation phase continued until Day 200.

### Infection of Retinal Organoids with AAV

Retinal organoids at Day 100 were placed individually in wells of a 96-well plate containing 50 µL of ProB27 medium. Organoids were transduced once with 4 × 10^9 vector genomes (vg) per organoid by directly adding the AAV vector to the culture medium. The AAV2-7m8 capsid was used to deliver the enhanced green fluorescent protein (EGFP) reporter gene under the control of one of the studied promoters.

Twenty-four hours post-infection, 50 µL of fresh ProB27 medium was added to each well and forty-eight hours after infection, organoids were washed twice with fresh ProB27 medium to remove residual viral particles and transferred to ultra-low attachment 24-well plates in DMEM:F12 (1:1, L-glutamine), 1% MEM non-essential amino acids, containing 2% B27 Supplement without vitamin A, 1 mM Glutamax, 40 U/mL penicillin, 40 µg/mL streptomycin, and 1 µg/mL amphotericin B. For each promoter, approximately 10 organoids were transduced per experiment, resulting in a total of around 40 organoids per promoter across four independent experiments.

Before stopping for quantification at D150, retinal organoids were examined using an Olympus inverted epifluorescence microscope to assess the presence of well-defined nuclear layers with maturing photoreceptor presenting outer segments^20^, and the level of GFP expression. Only organoids showing uniform lamination and outer segment development throughout their structure were retained for further analysis. For each promoter, organoids with similar levels of GFP expression were selected to ensure consistency across experimental conditions.

### NHP Retinal Explants Culture and infection with AAV

Eyes from a donated Cynomolgus macaque were removed and transported in cold CO2-independent medium (Thermo Fisher Scientific). The anterior segment and vitreous humor were removed, the retina was mechanically separated from the RPE and 5 mm punches were created in the peripheral retina with a sterile biopsy punch (Dutscher). The retinal explants were transferred onto polycarbonate transwell inserts (Corning) and cultured individually with Neurobasal-A + L-glutamine (final 2mM) plus B27 medium (Thermo Fisher Scientific). Two to 4 hours post-dissection, 1 µl of undiluted AAV was added on top of the retina explants.

Explants were cultured for two weeks in Neurobasal-A + L-glutamine (final 2mM) + B27 media at 37 °C in a 5% CO₂/95% air atmosphere.

### Quantification of GFP Expression by ddPCR

Laminated retinal organoids were collected at Day 150, gently dried, and stored at - 80°C until RNA extraction. Total RNA was extracted from individual organoids using the Quick-DNA/RNA Microprep Plus Kit (Zymo Research), following the manufacturer’s protocol. RNA concentrations were measured using a NanoDrop spectrophotometer (Thermo Fisher Scientific) to ensure sample quality and consistency. For reverse transcription, complementary DNA (cDNA) was synthesized using the SuperScript IV Reverse Transcriptase Kit (Thermo Fisher Scientific), according to the manufacturer’s instructions. Equal amounts of total RNA were used in each reaction to maintain consistency across samples. Digital droplet PCR (ddPCR) was performed using the QX200 Droplet Digital PCR System (Bio-Rad). Primers specific to GFP were used to quantify transgene expression, while GAPDH served as a housekeeping gene for normalization. ddPCR reactions were performed in technical triplicates to ensure data reliability. GFP expression levels were calculated by normalizing GFP copy numbers to GAPDH expression.

### Quantification of GFP Expression by Flow Cytometry Analysis

Retinal organoids at 50 days post infection were dissociated into single cells using papain (2 U, Worthington). The cell suspension was treated with DNase I (Sigma-Aldrich) to prevent clumping caused by released genomic DNA. Remaining large aggregates were removed by filtration, and the resulting single-cell suspension was stained with a viability dye (Miltenyi). Cells were subsequently fixed with 4% paraformaldehyde (PFA) at room temperature and permeabilized using saponin (eBioscience). Following permeabilization, cells were incubated with an anti-GFP antibody (BioLegend #338008) to enhance GFP detection. Stained samples were analyzed using BD FACSCelesta™ Flow Cytometer, and data were processed with FlowJo (10.9) software for downstream analysis.

### Immunostaining and Imaging

Mouse eyes, NHP retinal explants and retinal organoids were fixed in 4% paraformaldehyde (PFA) at room temperature for 2 h, 1h and 10 min, respectively. After fixation, samples were cryoprotected by immersion in PBS containing 30% sucrose overnight at 4 °C. Mouse eye cups were embedded in OCT compound and frozen in liquid nitrogen, while retinal organoids and NHP retinal explants were embedded in 7.5% gelatin and 10% sucrose in PBS, and frozen in dry ice-cold isopentane. Transverse cryosections were obtained using a cryostat (12 µm for mouse eyes, 10 µm for retinal organoids and NHP retinal explants). Sections were incubated in blocking solution (PBS, 1% BSA, 0.1% Tween-20, 0.1% Triton X-100 for mouse eyes; PBS, 0.2% gelatin, 0.1% Triton X-100 for retinal organoids and NHP retinal explants) for 1 h at room temperature. Primary antibodies diluted in the respective blocking solutions were applied overnight at 4 °C. The following primary antibodies were used: human CRX (Abnova, #H00001406-M02; 1:1000), human arrestin 3 (Novus Biologicals, #NBP1-37003; 1:500), mouse cone arrestin (Merck, #AB15282; 1:800), and GFP (AVES Labs, #GFP-1020; 1:2000). After PBS washes, sections were incubated with Alexa Fluor-conjugated secondary antibodies (1:600) for 2 h at room temperature, followed by DAPI nuclear staining. Mounted sections were visualized using a confocal microscope, and images were processed using ImageJ/FIJI software. Z-sections were projected into a single plane using maximum intensity projection.

### Quantification of GFP Expression by Automatic High Content Analysis

GFP expression was quantified using The CellVoyager CQ1 High-Content Analysis system (Yokogawa) for high-content imaging. Retinal sections were imaged using the Cellvoyager CQ1 spinning disk with a 20× objective. For each retinal slide, 20 fields were acquired as z-stacks spanning 13 µm with a step size of 1 µm.

On the CellPathFinder™ software, the quantitative analysis was performed on two-dimensional tile images generated from maximum intensity projections (MIP) of the acquired z-stacks. Regions of interest (ROIs) were defined within the nuclear layers. Nuclei were detected based on DAPI staining, and the resulting detection masks were applied to the other channels (GFP and arrestin). Positive cells were quantified using size- and intensity-based filters to exclude debris and background signals. Machine learning–based sorting was applied to improve the detection of cone arrestin–positive cells.

### Network of Transcription Factors

To discover transcription factors binding site motifs we used FIMO^22^. FIMO maps transcription factors binding sites onto a input sequence by comparing it to a database of known motifs corresponding to curated transcription factors binding sites. For this comparison we used the JASPAR core vertebrates non redundant dataset^23^. More precisely, this comparison is done through a letter-probability matrix, which, for a given motif, associates the probability of each nucleotide at a position in the motif’s sequence. To make the analysis more robust we selected the motifs with a q-value ≤ 0.05, following the standard threshold for p-values. The q-value corresponds to the positive false discovery rate. To obtain an understanding of the network of interactions between transcription factors both within a promoter (intra) and between promoters (inter), we first, counted and mapped the common binding sites both intra- and inter-promoters. This with the hypothesis that the local increase of density of transcription factors would benefit both sequences in term of rate of transcription. Second, we counted and mapped the transcription factors for which an interaction was documented. That is, transcription factors both present on the sequences-either on the same promoter (intra) or different promoters (inter)- for which there is experimental evidence of a physical interaction. For this last part we use the stringdb database (https://string-db.org). It should be noted that all the binding sites for the same transcription factor presenting an overlap of one or more nucleotides were removed from the analysis.

### Promoter Prediction and Fusion Analysis

Promoter activity was analyzed using a transformer-based DNA sequence classifier fine-tuned for promoter detection. The model backbone was the Nucleotide Transformer 500M–1000G^24^, and fine-tuning was performed using the official DNABERT-2 training code on the Genome Understanding Evaluation (GUE) promoter dataset, without modification^25^. The Nucleotide Transformer framework provides contextual DNA sequence representations suitable for regulatory element prediction^24^. To generate promoter probability profiles, each DNA sequence was scanned using overlapping 50 bp windows with a step size of 1 bp. For each window, the fine-tuned model produced a promoter probability score. Window-level scores were projected to base-resolution promoter probabilities by averaging the scores of all overlapping windows covering each nucleotide position. Predicted promoter regions were defined as contiguous intervals with promoter probability ≥ 0.5 and minimum length of 50 bp. Two fusion constructs were analyzed: Nocchu or GRK1–PR1.7 and Pikali PR1.7–GRK1, generated by concatenating the corresponding sequences in opposite orientations. Fusion junctions were explicitly annotated to assess how promoter activity redistributes across the fusion boundary. Promoter probability profiles were visualized as continuous curves with shaded high-probability regions.

### Statistical Analysis

Statistical analyses were performed using GraphPad Prism. An unpaired parametric t-test with Welch’s correction was used to compare the GFP expression between groups. No correction for multiple comparisons was applied, as the statistical tests were targeted to address specific hypotheses. Data are presented as mean ± SEM, with significance defined as p < 0.05. Each experiment was conducted with at least three biological replicates.

## RESULTS

### Promoter fusion for Combined Rod and Cone Expression

The design of promoters capable of targeting both rods and cones is a critical challenge in gene therapy for inherited retinal dystrophies. Rather than engaging in a labor-intensive search for de novo promoters, we hypothesized that combining two well-characterized and widely used promoters -one preferentially expression in rods and the other in cones- could generate a novel synthetic promoter capable of driving expression in both photoreceptors (Fig.1).

**Figure 1:**
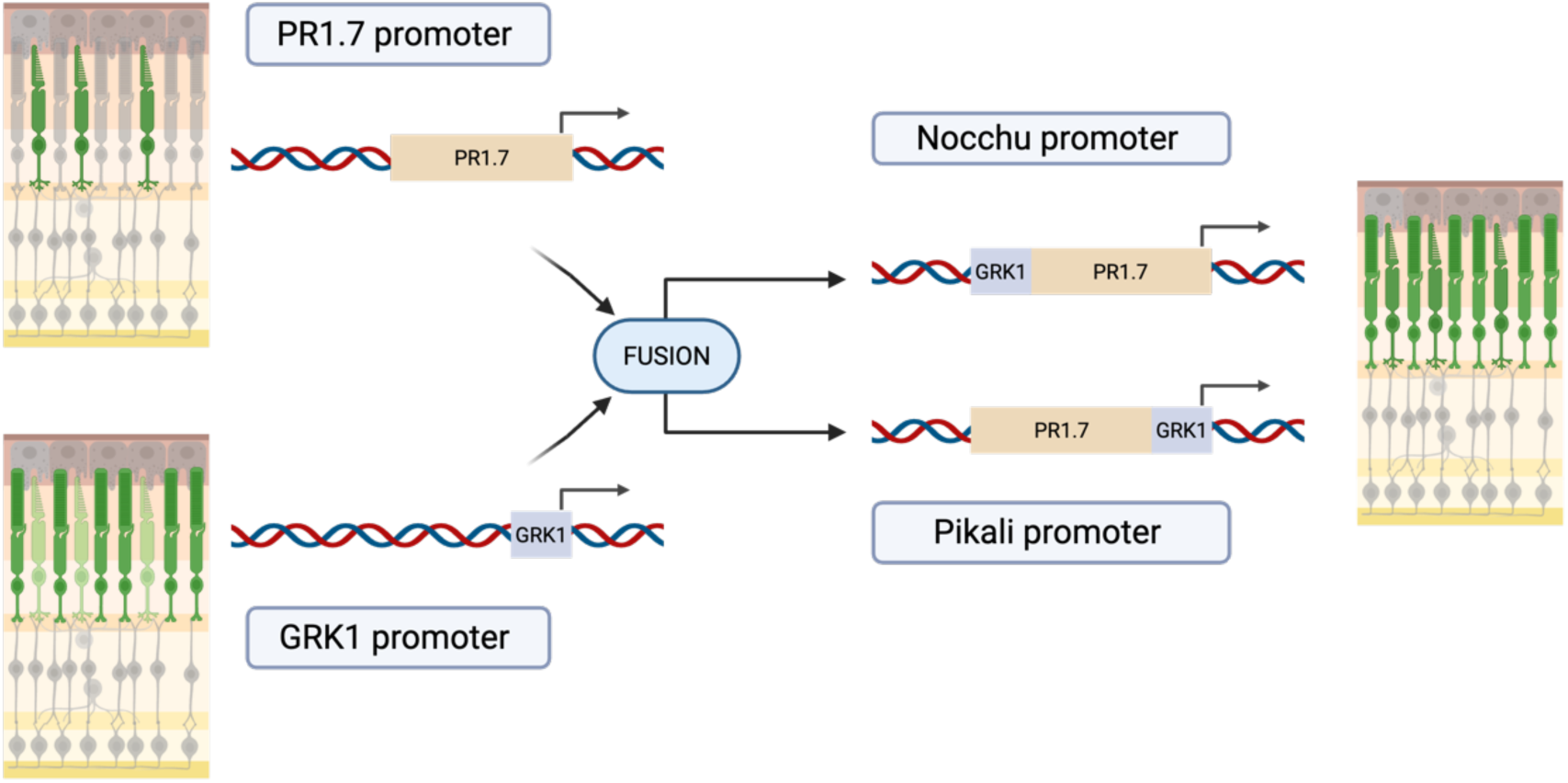
Design and Strategy for Fusion Promoters Targeting Rods and Cones. Schematic representation of the promoter design strategy. The cone-specific promoter PR 1.7 and rod-specific promoter GRK1 were fused in two configurations, creating the fusion promoters Nocchu and Pikali. These constructs aim to combine the specificity and expression strengths of their parental promoters to achieve dual targeting of rods and cones. The fusion promoters were tested for their ability to drive gene expression across both photoreceptor subtypes, providing a balanced and efficient tool for retinal gene therapy.

As a cone-specific promoter, we selected the opsin promoter PR1.7, known for its strong expression in cones and extensively studied in our laboratory^3^. To complement this, we chose the rhodopsin kinase promoter GRK1 as a rod promoter due to its small size of approximately 300 bp^6^. Given the packaging capacity limit of approximately 4.7 kb for AAV vectors, minimizing the size of the promoter is critical to allow sufficient space for the therapeutic gene and other necessary regulatory elements. Although the rhodopsin promoter is widely used for rod-specific targeting, GRK1 was favored here because it has the potential to drive expression in both rods and cones^5,6^, making it more suitable for a pan-photoreceptor strategy.

To create our promoters, we fused these two sequences in both configurations, resulting in Pikali promoter when PR1.7 was placed upstream of GRK1, and Nocchu when GRK1 was placed upstream of PR1.7. This approach generated two synthetic promoters of approximately 2 kb each, designed to integrate the strengths of their parental elements, with the expectation of achieving efficient targeting of both rods and cones (Fig.1).

Thus, our study focused on four promoters: the fusion promoters Pikali and Nocchu, alongside the original PR1.7 and GRK1 promoters used as benchmark promoters. This design allowed us to systematically assess the efficiency, specificity, and expression patterns of each promoter in both *in vitro* and *in vivo* models.

### Validation of Fusion Promoter Activity in the Mouse Retina

Following the design of the fusion promoters, we searched to validate their functionality and specificity in a well-established model for preclinical evaluation of retinal gene therapies like the mouse retina. This step was critical to confirm that the promoters could drive efficient and targeted expression in photoreceptor cells, laying the groundwork for their potential applications. In our analysis, GFP was used as a reporter gene to evaluate both cell-type specificity and signal intensity. Immunostaining on mouse retinal cryosections revealed distinct patterns of expression across the ONL, reflecting variations in both localization and intensity depending on the promoter.

Across all four promoters, GFP fluorescence was consistently restricted to the outer nuclear layer (ONL), a region anatomically exclusive to photoreceptors, confirming the photoreceptor-specific activity of these promoters (Fig.2C, F, I, L). For the benchmark promoters GRK1 and PR1.7, distinct expression patterns were observed. The PR1.7 promoter (Fig.2C) exhibited bright GFP expression in cones, as evidenced by colocalization with cone arrestin (CAR), with an expression in rods, which has been previously observed in the murine retina^3,4^. In contrast, the GRK1 promoter displayed expression in rods, with no detectable GFP signal in cones, as shown by the absence of colocalization between GFP and CAR (Fig.2F).

**Figure 2.**
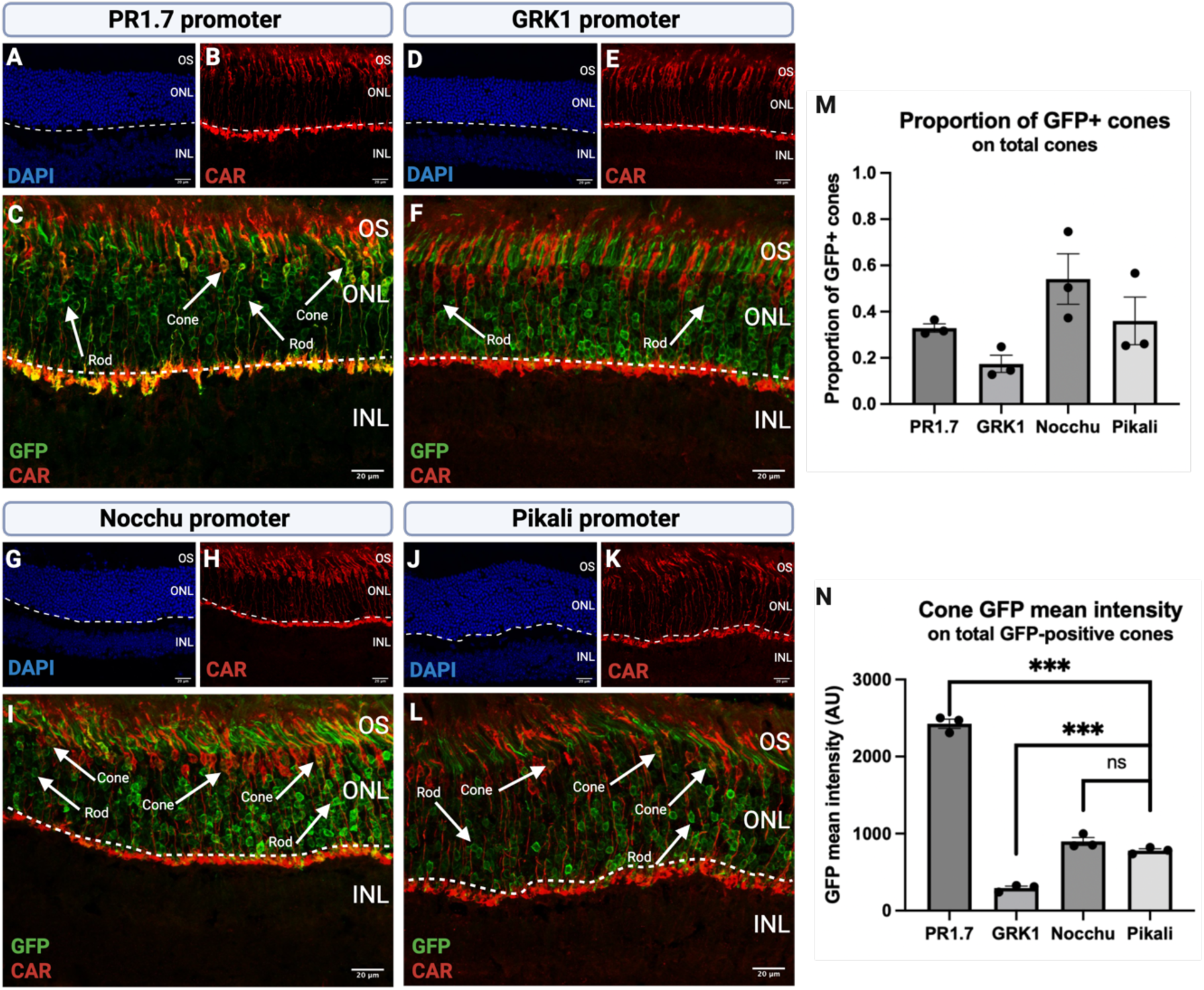
Immunostaining and quantification of promoter-driven GFP expression in mouse photoreceptors. Each set of panels corresponds to a specific promoter: (A-C) PR1.7, (D-F) GRK1, (G-I) Nocchu, and (J-L) Pikali. DAPI staining (A, D, G, J) highlights retinal organization, with dotted lines marking the boundary between the ONL and INL. Cone arrestin (CAR) staining (B, E, H, K) identifies cone photoreceptors, while merged images of CAR and GFP staining (C, F, I, L) reveal promoter-specific GFP expression and colocalization in photoreceptors. Arrows indicate GFP positive cones and rods. Confocal images were acquired using a 60 x oil objective. Scale bars: 20 pm. ONL, outer nuclear layer; INL, inner nuclear layer; OS, photoreceptor outer segments. (M) Quantification of GFP-positive cones relative to the total cone population. (N) Mean GFP fluorescence intensity in GFP-positive cones. GFP expression was quantified on maximum intensity projections of z-stack images acquired using a CQ1 Confocal Quantitative Image Cytometer. Each data point represents one eye (n = 3 eyes per condition). Data are presented as mean ± SEM. Statistical significance was determined using an unpaired parametric t-test with Welch’s correction (*p < 0.05, **p < 0.01, ***p < 0.001; ns, not significant).

Having established the cell-specific activity of the benchmark promoters GRK1 and PR1.7, we next evaluated the performance of the fusion promoters Pikali and Nocchu to determine whether they effectively integrate the strengths of their parental elements and achieve robust expression in both rods and cones. The Pikali and Nocchu promoters demonstrated broad expression across photoreceptors, with GFP detected in both rods and cones (Fig.2I, L). However, while expression in rods was strong and consistent, cone expression was less robust than anticipated. Although cone expression was less intense and observed in fewer cells compared to the PR1.7 promoter, it was notably improved relative to the GRK1 promoter. These results confirm the enhanced capacity of the Pikali and Nocchu fusion promoters to target both rods and cones in the mouse retina.

### Photoreceptor and Cone Transduction Efficiency in the Mouse Retina

Photoreceptor transduction efficiency in the mouse retina was quantified by automatic high-content analysis of GFP expression on retinal sections. Cone transduction was assessed by measuring both the proportion of GFP-positive cones, identified by cone-arrestin labeling, and the GFP mean fluorescence intensity within cone photoreceptors.

The proportion of GFP-positive cones relative to the total cone population differed between conditions. PR1.7 and Nocchu tended to show higher proportions of GFP-positive cones compared to GRK1, with Pikali displaying intermediate values; however, these differences were not statistically significant (Fig.2M).

Quantitative analysis of GFP mean intensity in cone photoreceptors revealed significant differences across conditions. PR1.7 resulted in the highest GFP signal in cones, while GRK1 exhibited substantially lower fluorescence intensity. Nocchu and Pikali showed intermediate levels, with no significant difference between these two promoters (Fig.2N).

Encouraged by these results, we next evaluated the performance of these promoters in retinal organoids derived from human iPSCs. This model, which closely mimics the gene expression profile of the human retina, provides a critical step toward assessing the clinical translation potential of these promoter constructs.

### Validation of Promoter Activity in Human Retinal Organoids

To evaluate the efficiency of the studied promoters in driving GFP expression, retinal organoids were infected with AAV2-7m8 vectors at D100 and analyzed at D150.

Representative images of organoids in culture demonstrate strong GFP fluorescence for the Pikali, Nocchu, and PR1.7 promoters, whereas the GRK1 promoter showed a markedly reduced signal. Non-transduced (NT) controls confirmed the absence of background fluorescence. These images were captured under identical acquisition settings, ensuring a fair comparison across conditions (Fig.3A).

**Figure 3:**
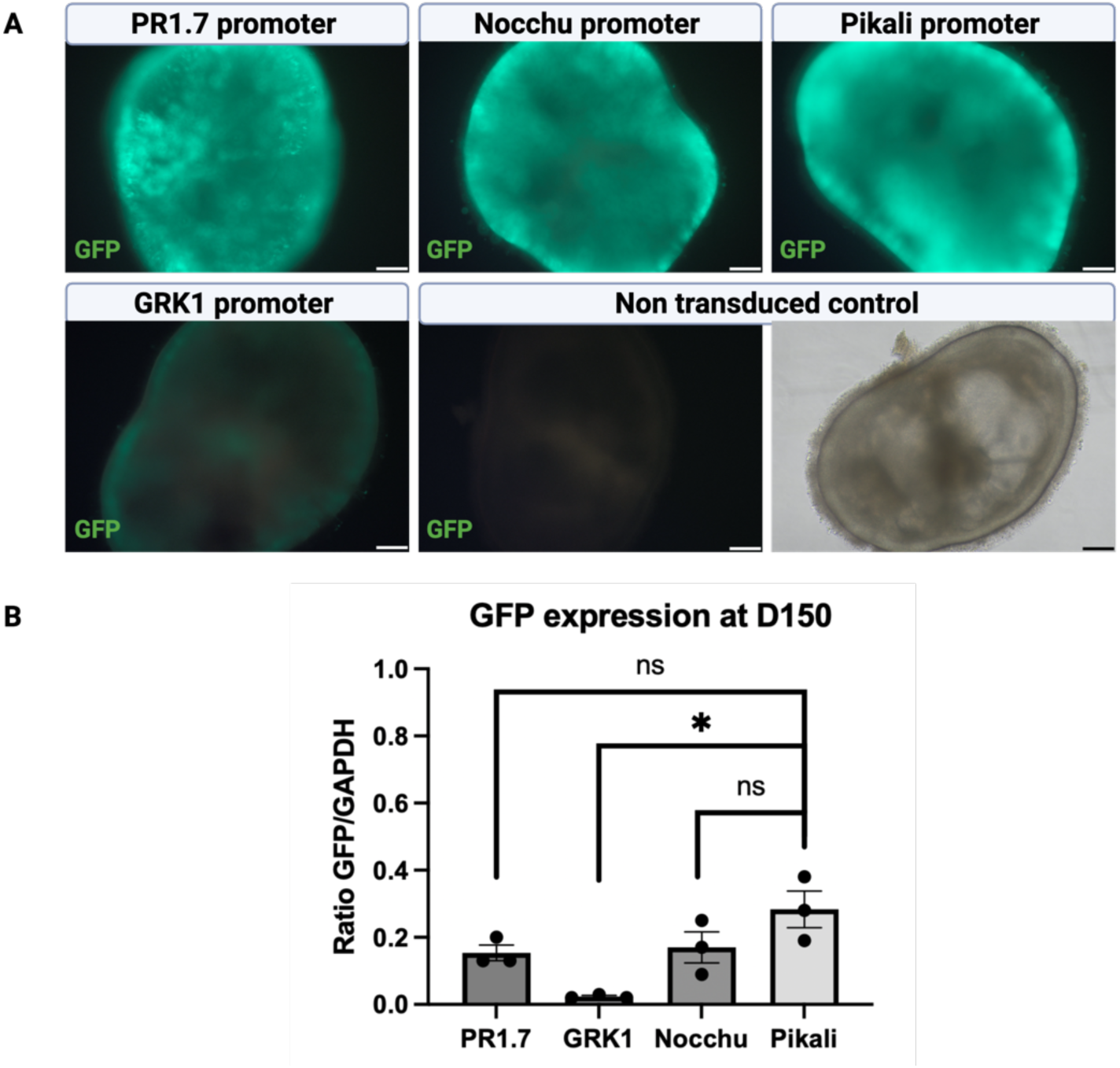
Comparison of GFP expression under different promoters in retinal organoids at D1S0. (A) Representative images of retinal organoids at 50 days post-infection (D150). GFP signal was captured using identical imaging parameters to ensure comparability across promoters. Brightfield imaging (shown only for the non-transduced control) was used to select organoids with a similar laminated structure and outer segments, indicative of proper development post-infection. Images were acquired using epifluorescence microscopy at 10x magnification. Scale bar: 100 pm. (B) GFP expression levels normalized to the housekeeping gene GAPDH, measured by ddPCR. Data are presented as mean ± SEM and analyzed using an unpaired parametric t-test with Welch’s correction. Expression levels were quantified at DI50, using organoids with a laminated structure consistent with the non-transduced control.

Quantitative analysis of GFP mRNA levels by RT-ddPCR further corroborated the visual observations. The Pikali promoter achieved the highest expression levels, followed by Nocchu and PR1.7, which displayed comparable but slightly reduced activity. In contrast, the GRK1 promoter showed significantly lower expression. Statistical analysis revealed significant differences between Pikali and GRK1 (p < 0.05), highlighting the superior transcriptional efficiency of Pikali compared to GRK1 (Fig.3B).

Together, these results confirm the robust performance of Pikali and Nocchu fusion promoters in driving GFP expression in retinal organoids.

### Cell-Type Specificity of Promoter Activity in Retinal Organoids

To evaluate the cellular specificity of GFP expression driven by the tested promoters, immunostaining was performed on retinal organoid cryosections (Fig.4). Photoreceptors were labeled with CRX, cones were identified by ARR3. Across all conditions, GFP expression was restricted to CRX-positive cells, confirming that all tested promoters specifically target photoreceptors. However, the localization and intensity of GFP expression within photoreceptors differed among the tested promoters. Consistent with observations in mice, the GRK1 promoter demonstrated strict specificity for rods, with GFP fluorescence exclusively detected in CRX-positive, ARR3-negative cells (Fig.4F). No GFP signal was observed in cones, further emphasizing the rod-specific nature of this promoter, even in human tissue. In contrast, the PR1.7 promoter exhibited a GFP expression limited to sparse CRX- and ARR3-positive cells, indicating highly restricted activity within a small subset of cones (Fig.4C). For the Nocchu promoter, GFP expression was predominantly observed in cones, but a subset of rods also exhibited weak GFP fluorescence (Fig.4I). In contrast, the Pikali promoter exhibited robust GFP expression in both cones and rods, with strong fluorescence detected in CRX-positive cells regardless of ARR3 expression (Fig.4L).

**Figure 4:**
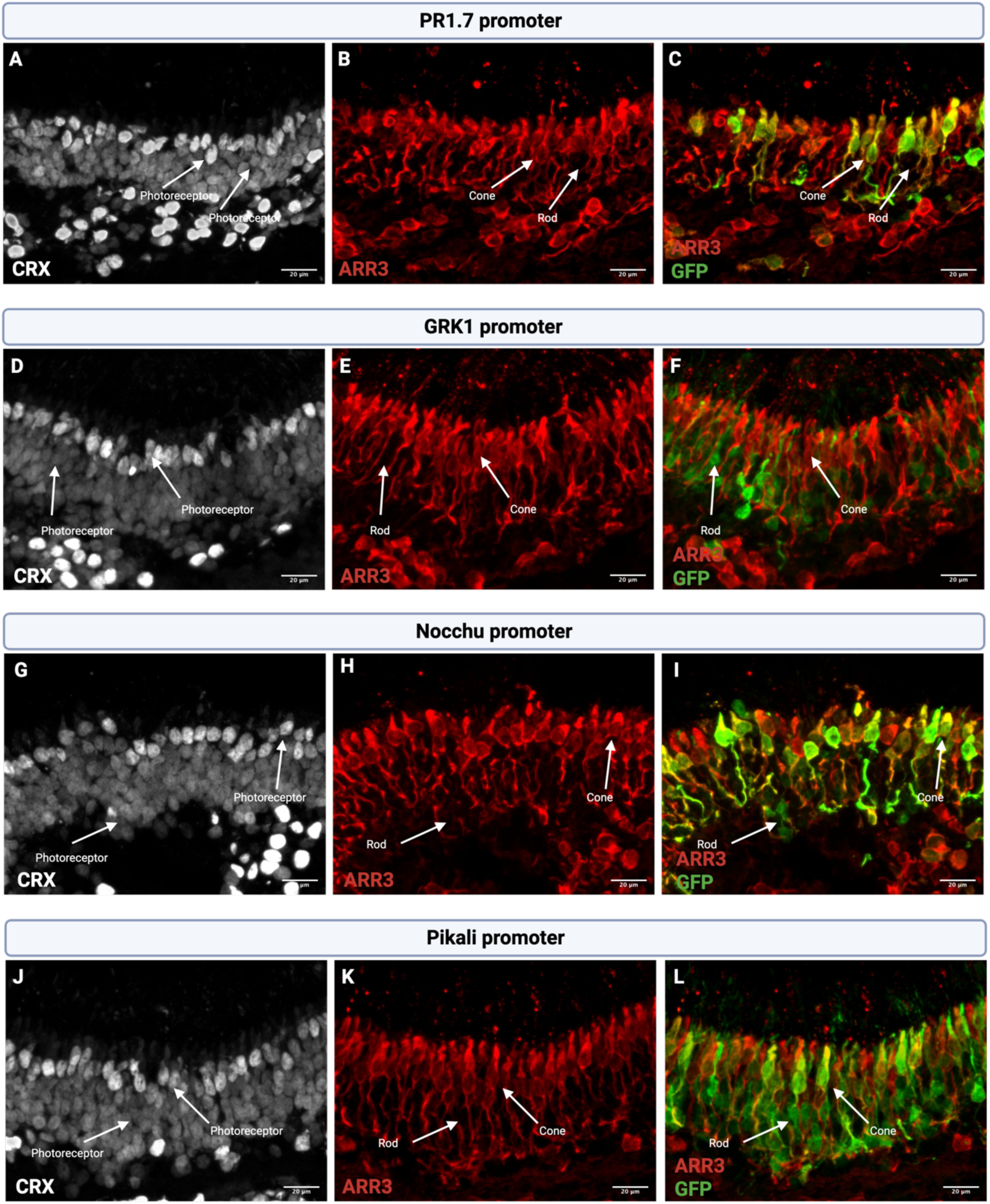
Immunostaining of human retinal organoid sections to assess cell-type specificity of promoter activity. Each set of panels corresponds to a specific promoter: (A-C) PR 1.7 promoter, (D-F) GRK1 promoter, (G-I) Nocchu promoter, and (J-L) Pikali promoter. Within each set, CRX staining (A, D, G, J) marks photoreceptors, ARR3 staining (B, E, H, K) distinguishes cones (CRX^+^ARR3^+^) from rods (CRX^+^ARR3“), merged images of ARR3 and GFP staining (C, F, I, L) shows promoter-driven GFP expression in specific photoreceptor cell types. Confocal images were acquired with a 40* oil-immersion objective ana 2x digital magnification. Scale bar: 20 pm.

These results suggest that the fusion promoters are promising, as they successfully drive GFP expression in both rods and cones. However, a more in-depth analysis is required to determine which promoters achieve the highest efficiency and specificity in targeting photoreceptors.

### Photoreceptor and Cone Transduction Efficiency in Retinal Organoids

To evaluate the transduction efficiency of our promoters in retinal organoids, we used two complementary fluorescent reporter cell lines. In the first, all photoreceptors expressed mCherry under the control of the CRX promoter, AAVS1::CrxP_H2BmCherry enabling the identification of photoreceptors. The graph showing the percentage of photoreceptors transduced for each promoter, highlights the differences in expression efficiency among the tested constructs (Fig.5B). Among the tested promoters, Pikali achieved the highest transduction rate, with approximately 45% of photoreceptors GFP-positive. Nocchu followed with about 30%, while the GRK1 promoter showed weaker transduction efficiency, with just over 20% of photoreceptors GFP-positive. In contrast, the PR1.7 promoter exhibited minimal efficiency, transducing only a very small fraction of photoreceptors.

**Figure 5:**
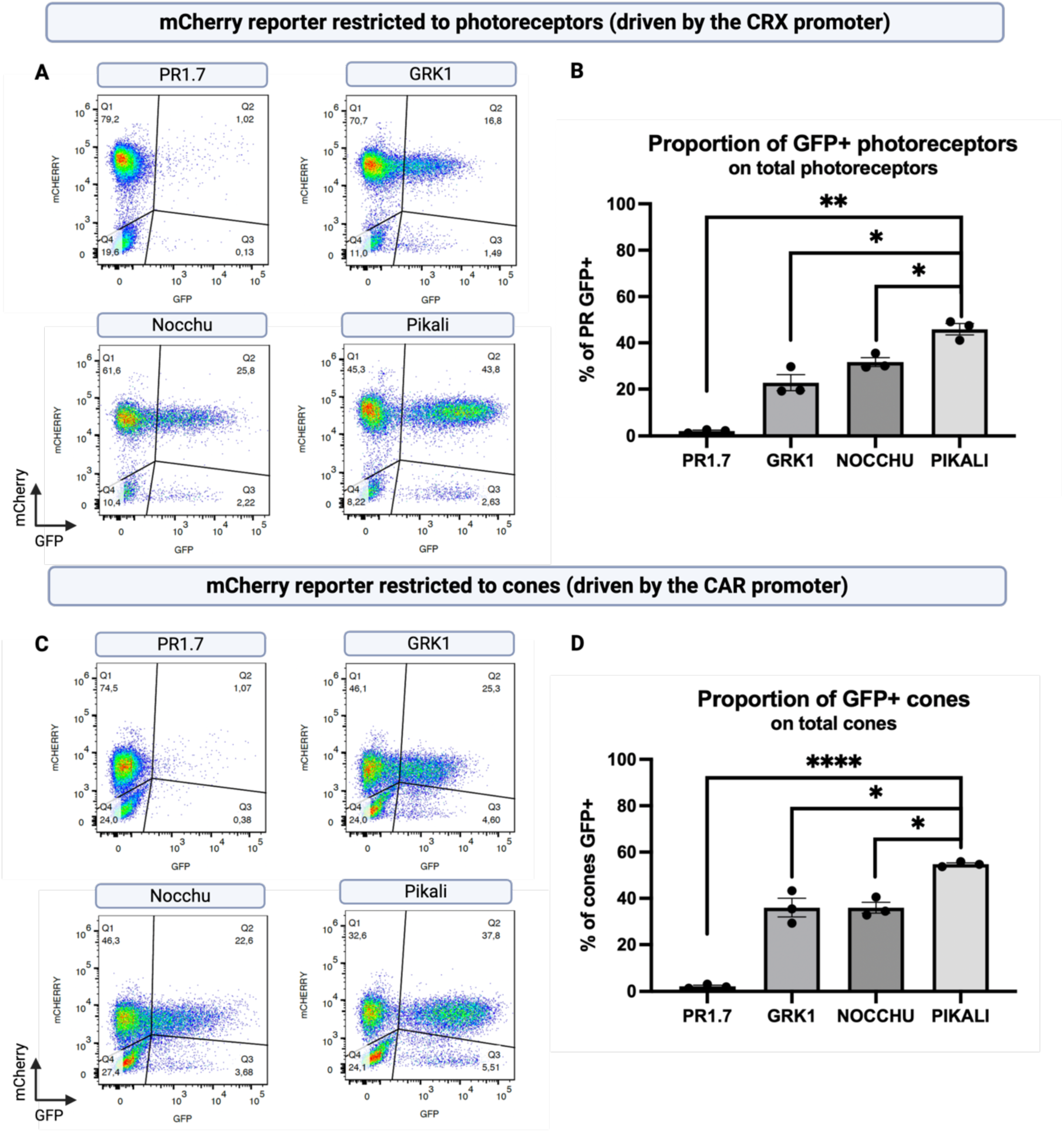
Quantification of GFP^+^ photoreceptors and cones in retinal organoids by flow cytometry. (A) Representative dot plots showing GFP expression under different promoters in photoreceptors (PRs) at DI50. The mCherry marker identifies the photoreceptor population (Q1 + Q2), with GFP^+^ photoreceptors localized in Q2. (B) Proportion of GFP^+^ photoreceptors within the total photoreceptor population, calculated as percentage of GFP^+^ photoreceptor in Q2 over total photoreceptors (Q1 + Q2). (C) Representative dot plots showing GFP expression under different promoters in cones at DI50. The mCherry marker identifies the cone population (Q1 + Q2), with GFP^+^ cones localized in Q2. (D) Proportion of GFP^+^ cones within the total cone population, calculated as percentage of GFP^+^ cones in Q2 over total cones (Q1 + Q2). (B and D) Data represent mean ± SEM from three independent experiments. Statistical significance was determined using an unpaired parametric t-test with Welch’s correction (*p < 0.05, **p < 0.01, ****p < 0.0001).

Representative dot plots from independent experiments illustrate the distinct GFP expression profiles for each promoter, providing additional insights into the levels of GFP expression within photoreceptors (Fig.5A). The Pikali promoter demonstrated a strong and uniform GFP signal across the transduced photoreceptor population, while the Nocchu promoter showed intermediate GFP intensity. The GRK1 promoter exhibited weaker GFP expression, with most signals clustering near the negativity threshold. Interestingly, the PR1.7 promoter displayed very high GFP intensity, but only in a few photoreceptors, suggesting a highly restricted but strong activity within a limited subset of cells.

To refine these observations, we performed a second analysis using the AAVS1::Arr3P_H2BmCherry reporter line (Fig. S2). In this line, mCherry expression was restricted to a subset of CRX+ photoreceptors and completely overlapped with CAR staining, confirming its cone specificity. This enabled specific evaluation of GFP expression in cones (Fig. 5C, D). The results mirrored the trends observed across the entire photoreceptor population, with the Pikali promoter demonstrating the highest efficiency, transducing 55% of cones. The Nocchu and GRK1 promoters both transduced approximately 35% of cones.

However, while Nocchu-driven GFP expression was easily detectable by immunostaining, GRK1-driven expression remained below the detection threshold, supporting substantially lower expression levels in cones. The PR1.7 promoter again showed very limited GFP expression, with only 2% of ARR3-mCherry-positive cones expressing GFP despite more than 60% of photoreceptors in the organoids being ARR3-mCherry positive, indicating a high proportion of cones. This further supports its highly restricted activity.

Together, the data highlight the superior performance of the Pikali promoter in driving robust GFP expression across both photoreceptors and cones, with Nocchu providing intermediate efficiency. The GRK1 promoter achieved broader but weaker transduction, while the PR1.7 promoter demonstrated strong but highly restricted expression. These findings underscore the distinct characteristics of each promoter, raising important questions regarding their specificity and efficiency in the murine and human retina.

### Cell-Type Specificity of Promoter Activity in Retinal Explants

To extend our observations beyond murine and human organoid models, we evaluated AAV-mediated transduction in non-human primate (NHP) retinal explants maintained in culture for two weeks using the same AAV constructs. Overall, transgene expression remained limited across all constructs, consistent with the known challenges of achieving efficient transduction in primate photoreceptors under ex vivo conditions. This was particularly evident in cone photoreceptors, which are both less abundant and more vulnerable in culture, and may be further disadvantaged by competition with rod outer segments for AAV access. Despite the low overall signal, distinct promoter-dependent patterns emerged.

PR1.7-driven expression was sparse and appeared restricted to cones (Fig.6A-C), although transduction efficiency in this population was lower than expected based on previous reports in *in vivo* NHP studies and human retinal explants^3^. Nocchu exhibited similarly low overall expression levels but showed a modest increase in cone transduction compared to PR1.7, along with low but detectable expression in rods, resulting in a broader cellular distribution (Fig.6G-I). In contrast, Pikali drove strong and widespread expression in rods, with a large proportion of cells appearing GFP-positive across the imaged areas (Fig.6J-L). This high level of rod transduction made the identification of cone expression more challenging; however, occasional GFP-positive cones were detected, at a frequency that appeared closer to PR1.7 than to Nocchu. GRK1-driven expression was observed predominantly in rods, with no clear signal detected in cones in this dataset (Fig.6D-F); however, given the overall low cone transduction efficiency observed across constructs, this apparent absence should be interpreted with caution. Importantly, the relative promoter-specific patterns observed in NHP explants are consistent with those obtained in human retinal organoids, supporting the reproducibility of promoter behavior across complementary model systems despite differences in overall transduction efficiency.

**Figure 6:**
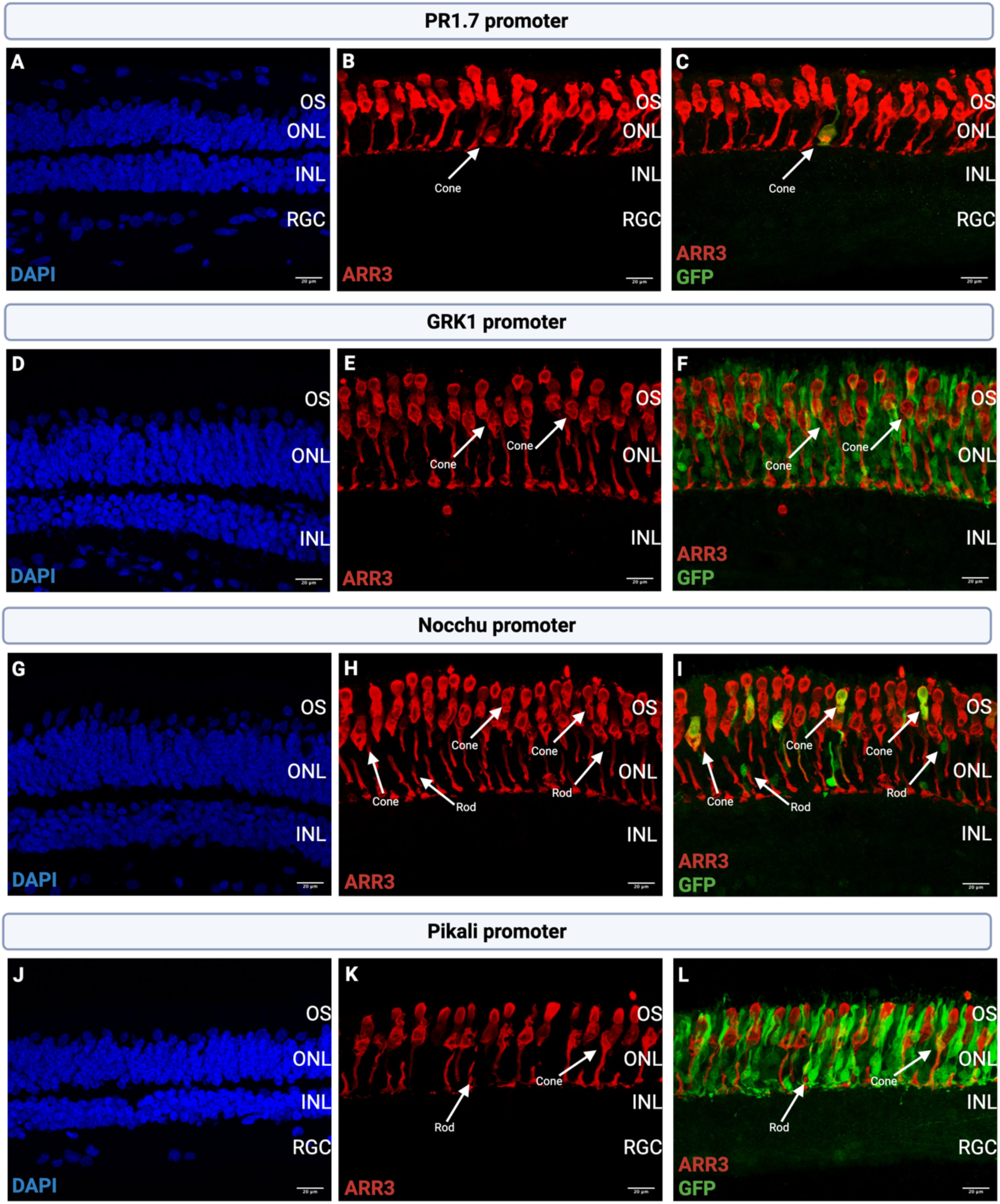
Immunostaining of NHP retinal explants to assess promoter-driven GFP expression in photoreceptors. Each set of panels corresponds to a specific promoter: (A-C) PR1.7 promoter, (D-F) GRK1 promoter, (G-I) Nocchu promoter, and (J-L) Pikali promoter. Within each set, DAPI staining (A, D, G, J) highlights retinal layers. Staining with cone arrestin, CAR, (B, E, H, K) identifies cone photoreceptors, and merged images of ARR3 and GFP staining (C, F, I, L) reveals promoter-specific expression patterns in photoreceptors. Arrows indicate GFP^+^ rods and cones for PR 1.7, Nocchu, and Pikali promoters, although cone transduction appears limited. In contrast, the GRK1 promoter (E, F) shows GFP expression restricted to rods, with no detectable signal in cones. Confocal images were acquired using a 60* oil objective. Scale bars: 20 pm. ONL, outer nuclear layer; INL, inner nuclear layer; OS, photoreceptor outer segments.

### Network of Transcription Factors

In order to understand the possible mechanisms beyond the improved performance of our fusion promoters we studied the transcription factors (TFs) and binding sites distribution across the two regulatory regions. Key assumption is that putative interaction between the TFs that bind to the two parental promoters can increase their own recruitment. To count inter-promoter putative interactions (Fig.7A) we consider the presence, in the two regulatory elements, of binding sites for either the same TF (Fig.7B) or pair of TF that are known to interact (Fig.7C and Methods). For both kinds, the number of putative interactions is largely increased by the fusion of the two parental promoters. These results show a densification in the interaction network for transcription factors between the two promoter sequences, i.e. the emergence of inter-promoters interactions in a scale that is either superior or comparable to the number of interactions on each individual promoter. This increased network connectivity could lead to a higher recruitment of transcription factors, thus explaining the increase in transcriptional activity in the fusion promoters (Fig.7A). However, this analysis does not differentiate between the two possible orientations.

**Figure 7:**
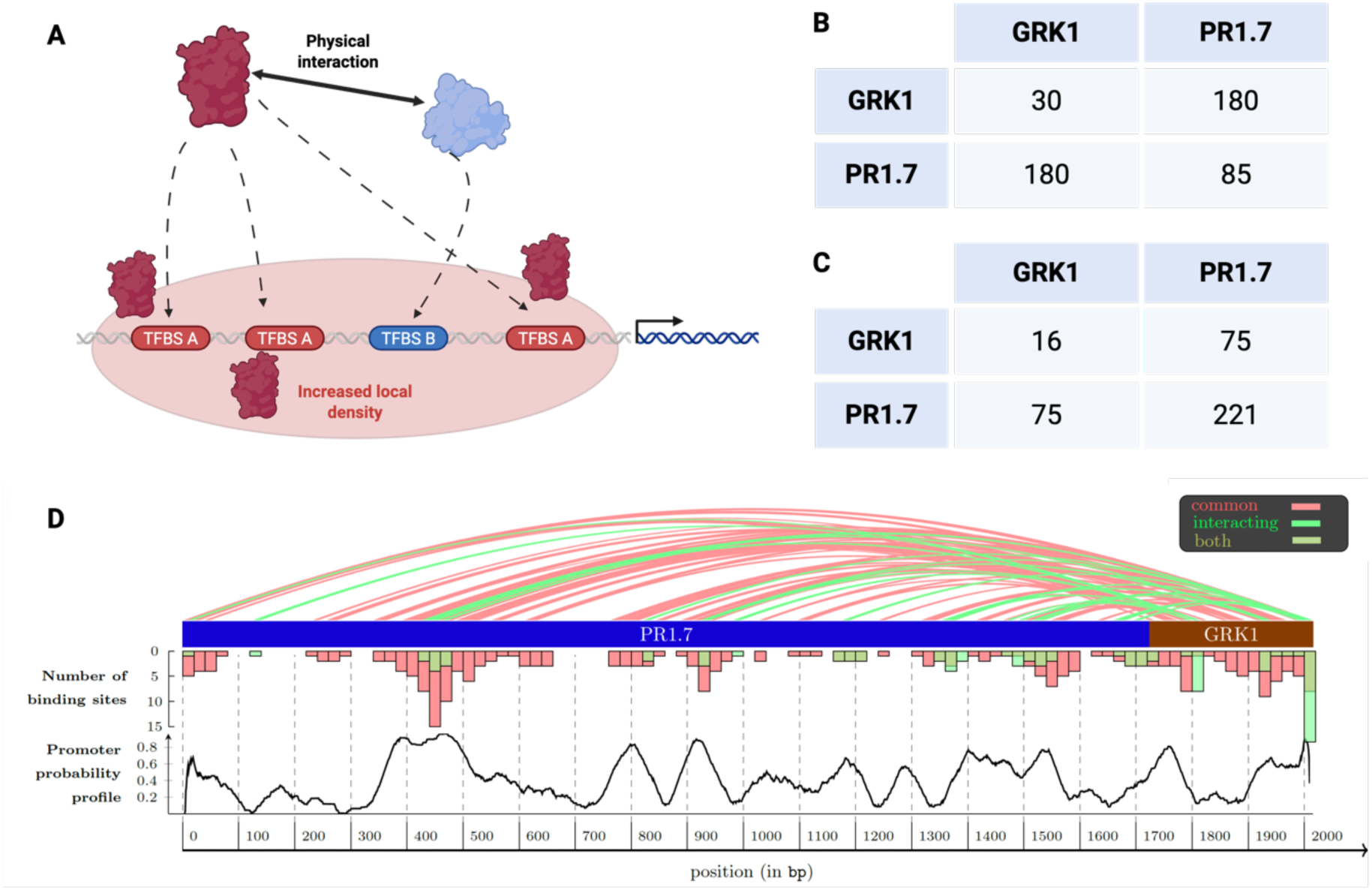
Transcription factor interaction network and regulatory architecture underlying promoter activity. (A) Schematic representation illustrating how increasing the number of transcription factor (TF) binding sites enhances TF recruitment, leading to a higher local TF density. This increased density promotes transcriptional activity and can further facilitate the recruitment of additional TFs through physical protein-protein interactions, thereby reinforcing promoter efficiency. (B) Number of identical transcription factor binding sites either within the same promoter sequence (diagonal elements) or shared between two promoter sequences (off-diagonal elements). (C) Number of interactions between transcription factors. Values correspond to the number of possible combinations of interacting TFs. Diagonal elements represent interactions between TFs whose binding sites are located within the same promoter sequence, whereas off-diagonal elements correspond to interactions between TFs whose binding sites are distributed across the two promoter sequences. (D) Distribution of transcription factor binding sites along the Pikali promoter shown as a histogram, together with the promoter activity probability profile (detailed in Supplementary Fig. 1). Arcs connecting pairs of binding sites represent either identical transcription factor binding sites shared between PR 1.7 and GRK1 sequences or pairs of binding sites corresponding to interacting transcription factors.

### Orientation-Dependent Regulatory Effects in Fusion Constructs

In order to understand the spatial organization of the regulatory region of our fusion promoters, and thus the difference between the two fusion order of Pikali and Nocchu, we applied a transformer-based DNA sequence classifier fine-tuned for promoter detection (Methods). The AI model is capable of identifying the sequence part that are likely to be involved in the regulation, and when applied to the two parental promoters it provided fundamentally different regulatory architectures. GRK1 exhibits a sharp, high-confidence promoter peak localized near a putative transcription start site, consistent with a compact core promoter optimized for transcription initiation (Fig. S1). In contrast, PR1.7 displays broader and more distributed promoter probability peaks, indicative of a more complex regulatory region with distributed promoter-like activity (Fig. S1). Fusion orientation strongly influences promoter behavior. In Pikali, the PR1.7→GRK1 construct (Fig. 7D and Fig.S1A), promoter probability accumulates continuously upstream of the GRK1 core promoter, producing a coherent region of elevated activity across the fusion junction. This configuration resembles a canonical regulatory architecture in which distributed regulatory elements precede and reinforce a compact core promoter^26^. In contrast, Nocchu, the GRK1→PR1.7 fusion exhibits a pronounced valley of low promoter probability around the fusion junction, separating the GRK1 core promoter from the downstream PR1.7 regulatory region (Fig. S1). This arrangement yields a fragmented promoter landscape and reduced continuity of promoter-like activity (Fig.S1B).

## DISCUSSION

In this study, we evaluated the transduction efficiency and cellular specificity of four promoters, PR1.7, GRK1, and their fusion constructs named Nocchu and Pikali, in both murine retinas and human retinal organoids. Our findings reveal distinct GFP expression patterns across photoreceptor subtypes, offering insights into their potential for gene therapies targeting both rods and cones.

Initial experiments in the mouse retina highlighted clear differences in cellular specificity and expression efficiency among the tested promoters. The GRK1 promoter drove expression predominantly in rods, with significantly lower GFP levels compared to PR1.7 in both rods and cones. Although previous studies have reported GRK1-driven expression in both photoreceptor subtypes, our results showed limited cone expression and reduced overall expression intensity. This discrepancy may be explained by differences in experimental conditions. For instance, Khani et al. used a higher AAV injection volume (2 µL at 1 × 10¹² vg/mL)^6^ compared to the 1 µL at the same titer used in this study. Higher vector doses have been associated with increased off-target transduction and may facilitate cone expression. In addition, the use of Nrl⁻/⁻ mice in prior studies, a cone-dominant model lacking rods, likely promoted GRK1-driven expression in cones due to the absence of rod competition for AAV uptake^6^. In contrast, PR1.7 drove markedly higher GFP expression levels in cones, consistent with previous reports showing strong cone-biased activity in the mouse retina. However, occasional rod expression was observed, a phenomenon previously described in murine models^3,4^. Together, these results underline the limitations of the two benchmark promoters in the mouse retina: GRK1 provides relatively weak expression, whereas PR1.7 drives markedly higher expression levels in photoreceptors in the mouse retina. The fusion promoters Nocchu and Pikali displayed intermediate expression profiles in the mouse retina. While they did not reach the expression intensity of PR1.7, they consistently outperformed GRK1 and showed activity in both rods and cones, supporting their evaluation in a human-relevant model.

Experiments conducted in human retinal organoids validated and extended these observations. Quantification by flow cytometry and ddPCR revealed marked differences between promoters in both transduction coverage and expression level. The GRK1 promoter exhibited broad but weak expression, targeting approximately 20% of photoreceptors with low GFP intensity. While this confirms its reliability for rod targeting, the limited expression level raises concerns regarding its therapeutic relevance in the human retina, where efficient cone expression is critical for visual acuity. In contrast, the PR1.7 promoter drove very high

GFP expression but in a highly restricted population, accounting for approximately 2% of photoreceptors. Flow cytometry thus revealed limited cellular coverage, whereas ddPCR analysis showed overall expression levels comparable to those obtained with the fusion promoters, indicating very strong expression on a per-cell basis. Such high expression confined to a small number of photoreceptors could potentially impose an increased transcriptional and translational burden on individual cells, which may be suboptimal in a therapeutic setting. The fusion promoters Nocchu and Pikali addressed these limitations by achieving a broader transduction profile, with GFP expression observed in both rods and cones and transduction of approximately 30–45% of photoreceptors. Although their expression intensity was lower than that achieved with PR1.7 on a per-cell basis, they reached comparable overall expression levels by distributing transgene expression across a substantially larger photoreceptor population. Importantly, both fusion promoters also outperformed GRK1 by combining higher expression levels with broader photoreceptor transduction, thereby supporting a more balanced expression profile.

Comparison between the mouse retina and human retinal organoids highlights notable species-specific differences in promoter behavior. In the mouse retina, PR1.7-driven expression was occasionally detected in rods, whereas this rod leakage was absent in human retinal organoids, suggesting differences in transcriptional regulation between species. These observations emphasize the limitations of extrapolating promoter performance from murine models to human systems^11^. While mouse studies remain essential for initial characterization, human retinal organoids provide a more relevant platform to evaluate promoter efficiency and specificity in a human context. In this human-relevant model, the fusion promoters Nocchu and Pikali demonstrated a favorable balance between cellular coverage and expression intensity, overcoming key limitations associated with GRK1 and PR1.7. Their performance in human retinal organoids supports their potential utility for gene therapy strategies requiring efficient targeting of both rods and cones.

To further refine this cross-species comparison, we next assessed AAV-mediated transduction in non-human primate (NHP) retinal explants, providing access to a structurally intact and mature retinal tissue in a species more closely related to humans. However, the use of retinal explants also introduces specific constraints related to tissue viability and culture configuration. Due to the limited survival of explants over time, AAVs were applied with the photoreceptor layer oriented toward the air–medium interface to maximize direct vector exposure. While this configuration facilitates viral access, it may also compromise photoreceptor preservation, particularly for cones, which are anatomically positioned above rods and may therefore be more vulnerable under these conditions.

While reduced viability likely affects all photoreceptors in explant culture, cone photoreceptors appear disproportionately impacted in terms of detectable transduction. This may reflect both their intrinsic vulnerability, but also the combined effects of cellular composition and photoreceptor maturity. In contrast to human retinal organoids, which typically contain a higher proportion of cones and relatively immature rod photoreceptors at the time of infection, NHP retinal explants from peripheral regions are dominated by rods and display fully developed outer segments. Previous work has shown that rod outer segment maturation can influence AAV-mediated transduction efficiency^27^, suggesting that mature rod structures may preferentially capture AAV particles and limit access to cones.

Together, differences in photoreceptor abundance and maturation state between models may contribute to the observed shift in expression patterns, with relatively enhanced cone expression in organoids and reduced cone transduction in explants. These observations may also have translational implications, as retinal composition is altered in degenerative conditions, where rod loss could modify AAV accessibility and potentially favor cone-targeted gene delivery.

In retinal organoids, more than 60% of photoreceptors were cones, yet only 2% of them expressed GFP under the control of PR1.7. Similarly, although cones were present within the imaged regions of NHP retinal explants, PR1.7-driven GFP expression remained sparse in this population. In contrast, Nocchu and Pikali consistently drove expression in both rod and cone photoreceptors across models. Together, these observations suggest that the limited PR1.7-driven expression may not solely reflect technical limitations in cone transduction, but could also reflect selective promoter activity within a subset of cone.

Interestingly, the fusion of GRK1 and PR1.7 into the Nocchu and Pikali constructs not only achieved broader cellular coverage but also increased overall expression levels. While GRK1 alone showed weak but detectable cone activity by flow cytometry, its contribution to the fusion constructs significantly enhanced cone expression. At the same time, PR1.7’s initially restricted cone expression expanded to a larger cone population while maintaining robust rod targeting. If the fusion were purely additive, GFP expression would likely remain limited to smaller subsets of photoreceptors, as observed with the individual GRK1 and PR1.7 promoters. However, the broader and more homogeneous expression profiles observed with Pikali and Nocchu point toward cooperative regulatory mechanisms that enhance transgene expression, resulting in both expanded cellular distribution and balanced intensity across rods and cones. While these experimental observations clearly argue against a simple additive model, they do not, on their own, explain the regulatory basis of this cooperative behavior.

This prompted us to investigate the transcriptional architecture of the fusion constructs in greater detail, with a particular focus on transcription factor networks, promoter organization, and the impact of fusion orientation on regulatory activity.

In silico analyses revealed a densification of transcription factor (TF) interaction networks and the emergence of inter-promoter interactions within the fusion constructs, indicating that promoter fusion generates a regulatory landscape with increased connectivity. Such integration may facilitate cooperative TF recruitment and stabilization of transcriptional complexes, providing a mechanistic basis for the increased transcriptional output observed experimentally.

In addition to transcription factor network interactions, the predicted regulatory architectures of the parental promoters suggest that spatial organization of regulatory elements may play a critical role in shaping fusion promoter activity. GRK1 is characterized by a compact, high-confidence core promoter, whereas PR1.7 exhibits a broader and more distributed regulatory profile. The fusion of these distinct architectures does not simply juxtapose two independent regulatory units but instead reshapes the spatial organization of regulatory elements. This reorganization creates a novel regulatory context in which distributed regulatory inputs can converge on a defined core promoter, reinforcing transcriptional activity through coordinated regulation. Importantly, promoter orientation emerged as a critical determinant of regulatory coherence. In the Pikali configuration (PR1.7→GRK1), distributed regulatory elements precede and reinforce the GRK1 core promoter, resulting in a continuous and structured promoter landscape. In contrast, the Nocchu configuration (GRK1→PR1.7) introduces a discontinuity at the fusion junction, yielding a fragmented regulatory architecture. These orientation-dependent effects underscore that promoter fusion is inherently directional and that regulatory ordering plays a decisive role in enabling synergistic interactions.

Taken together, these observations suggest that fusion promoters function through cooperative and architecture-dependent mechanisms rather than through simple additive effects. Beyond mechanistic interpretation, in silico analyses may also serve as a valuable tool to refine fusion promoter design, providing a framework to guide the rational selection and arrangement of regulatory elements prior to experimental testing.

Our findings further highlight that promoter performance is strongly influenced by the biological context in which regulatory activity is evaluated. While murine models remain essential for initial functional characterization, they present limitations in predicting promoter behavior in human systems. For example, the rod leakage observed with the PR1.7 promoter in the mouse retina was absent in human retinal organoids, underscoring species-specific differences in transcriptional regulation. These observations emphasize the importance of assessing promoter activity directly in human-derived retinal tissue to enhance translational relevance.

Extending this concept further, promoter performance may be most accurately evaluated within disease-relevant contexts. A recent study by Sudharsan et al. proposed a paradigm shift in promoter selection for retinal gene therapy by prioritizing regulatory elements that remain active or are upregulated during retinal degeneration, rather than relying solely on expression profiles in healthy tissue^28^. This strategy is particularly relevant for mid-to late-stage retinal degeneration, where transcriptional remodeling leads to the downregulation of many canonical photoreceptor promoters.

In this context, our fusion promoter strategy can be viewed as complementary to approaches aimed at identifying disease-responsive regulatory elements. While our work focused on combining established photoreceptor promoters to improve expression robustness and cellular coverage, the same fusion principle could be applied to promoters selected based on their sustained or upregulated activity during degeneration. In human retinal organoids, the fusion promoters Nocchu and Pikali demonstrated that promoter combination can enhance transgene expression beyond that achieved by individual elements. Together, these observations suggest that integrating disease-responsive promoters into fusion constructs may represent an additional avenue to further optimize transgene expression in degenerating retinal environments.

In conclusion, the robust performance of Nocchu and Pikali in human retinal organoids positions these fusion promoters as promising candidates for gene therapy targeting inherited retinal dystrophies. Disorders such as RDH12- and CEP290-associated inherited retinal dystrophies, which require efficient targeting of both rods and cones due to early photoreceptor involvement, could particularly benefit from promoters capable of balancing broad cellular coverage with controlled expression intensity^8,29^. By overcoming the limitations of their parental promoters, Nocchu and Pikali offer a viable solution for therapeutic strategies requiring simultaneous targeting of both photoreceptor subtypes.

Future studies should focus on validating these fusion promoters in vivo, with particular attention to long-term stability, safety, and therapeutic efficacy in preclinical models. More broadly, the strategy of promoter fusion could be extended to other gene therapy applications requiring coordinated targeting of multiple cell types or tissues. Diseases such as Pompe disease, which involve both muscle and liver, may benefit from tailored fusion promoters designed to enhance tissue coverage and therapeutic outcomes, highlighting the broader potential of this approach for next-generation gene therapy development.

## Acknowledgments

We are grateful to M. Cornebois, J. Degardin, M. Simonutti, and the staff of the Animal and Phenotyping facilities for their assistance. We thank S. Fouquet from the Imaging Facility for image acquisition and analysis. We thank M. Lechuga from the High-Throughput Screening (HTS) Facility for image acquisition and analysis performed on the CQ1. We thank C. Delarasse and the staff of the Cellular and Tissue Phenotyping Platform for their assistance with flow cytometry experiments. We thank V. Fradot for NHP eye dissection and explant preparation. We also thank G. Labernede for AAV production and titration. Finally, we thank E. Léger-Charnay for her valuable support and insightful advice throughout this study.

Conflicts of Interest

S.T., O.G. and D.D. hold a patent on Pikali and Nocchu promoter sequences. A.S.-B. and O.G. are inventors on patents on hiPSC retinal differentiation and on the use of hiPSC retinal derivatives to treat retinal degeneration, licensed to Gamut Cell Tx (Tenpoint Therapeutics). The remaining authors declare no competing interests.

## Author Contributions

S.T. contributed to the conception, design, execution, and analysis/interpretation of all experiments and writing of the manuscript. J.T. contributed to the bioinformatic analysis. E.A-Z. contributed to NHP infection and revision of manuscript. C.N. contributed to iPSC culture and histological processing. A.S-B. contributed to iPSC culture. T.V.M. contributed to the bioinformatic analysis and writing of the manuscript. L.R. contributed to flow cytometry acquisition and data analysis. A.P. contributed to CQ1 image acquisition and data analysis. M.D. contributed to NHP infection and explants culture. U.F. contributed to the bioinformatic analysis and revision of manuscript. O.G. contributed to the conception, design, and interpretation of experiments, manuscript writing, and financial support. D.D. contributed to the conception, design, and interpretation of experiments, manuscript writing, and financial support. All authors read and approved the final manuscript.

