## Supplementary figures for "*Promfusion*: a synthetic fusion promoter enabling enhanced and balanced photoreceptor transgene expression"

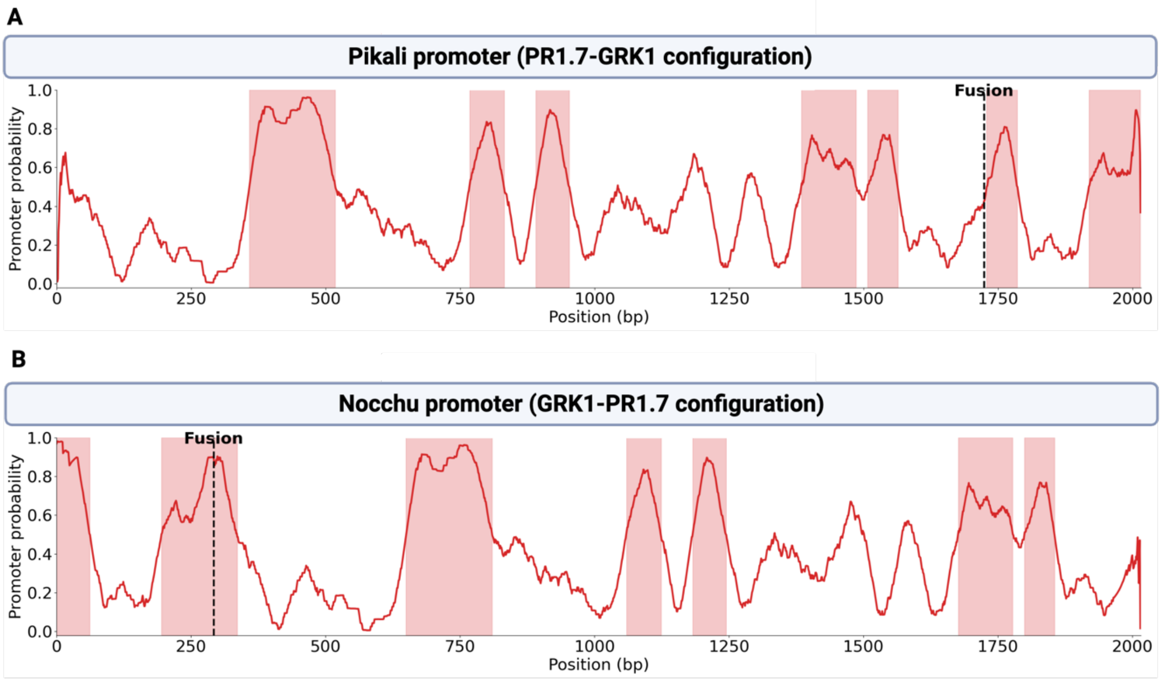


**Supplementary Figure 1. In silico promoter probability profiles of fusion constructs.** (A) Promoter probability along the full-length sequence of the Pikali fusion promoter (PR1.7–GRK1 configuration). (B) Promoter probability along the full-length sequence of the Nocchu fusion promoter (GRK1–PR1.7 configuration). Promoter probability is shown as a continuous curve along the sequence length (bp). Shaded regions indicate predicted promoter regions exceeding the probability threshold (≥ 0.5 ), and dashed vertical lines denote the fusion junction between the parental promoter sequences.

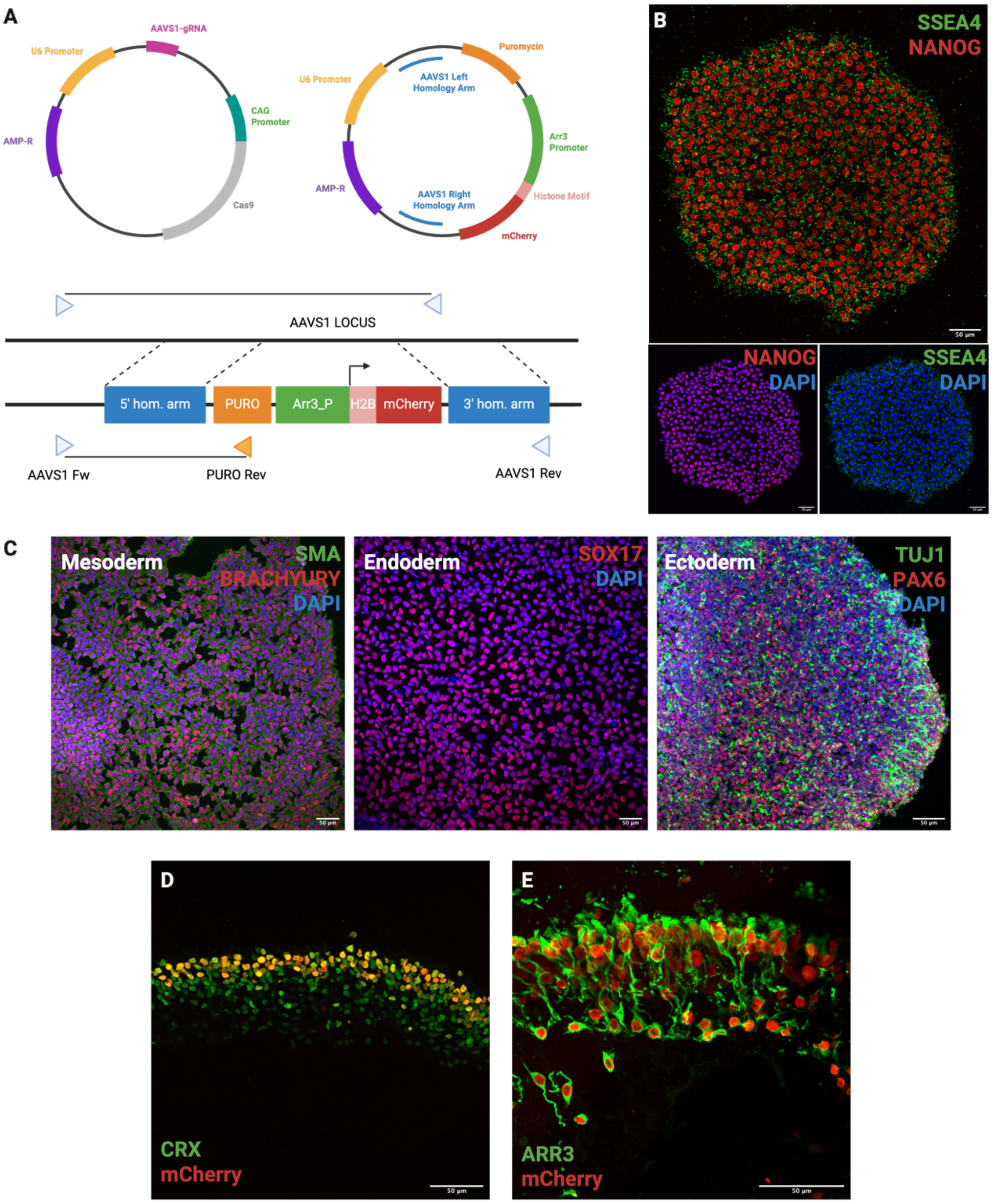


**Supplementary Figure 2. Generation and characterization of the AAVS1::Arr3P_H2BmCherry hiPSC line.** (A) Schematic representation of the CRISPR/Cas9 strategy and donor plasmid used to generate the AAVS1::Arr3P_H2BmCherry hiPSC line. Triangles indicate the positions of primer binding sites (AAVS1 Fw, PURO Rv, AAVS1 Rv) used to validate targeted integration of the fluorescent cassette. (B) Immunocytochemistry of pluripotency markers (NANOG and SSEA4) in AAVS1::Arr3P_H2BmCherry hiPSCs, confirming maintenance of pluripotency after genome editing. (C) Embryoid bodies (EBs) derived from AAVS1::Arr3P_H2BmCherry hiPSCs stained for markers of the three germ layers: endoderm (SOX17), mesoderm (BRACHYURY, SMA), and ectoderm (PAX6, TUJ1), demonstrating preserved differentiation potential. (D) Immunostaining for CRX in D120 retinal organoid sections showing that all mCherry⁺ nuclei co-localize with CRX⁺ photoreceptors, while a subset of CRX⁺ cells lack mCherry signal, consistent with the presence of non-cone photoreceptors. (E) Immunostaining for human cone arrestin 3 (ARR3) in D120 retinal organoid sections showing co-localization with mCherry⁺ cells, confirming that the reporter specifically labels cone photoreceptors.
